# Odon: An ultra-fast viewer for spatial proteomics

**DOI:** 10.64898/2026.03.30.715233

**Authors:** Alexander Coulton, Nicholas McGranahan

**Affiliations:** UCL Cancer Institute, Paul O’Gorman Building, 72 Huntley St, London WC1E 6DD

## Abstract

Multiplexed spatial proteomics and spatial transcriptomics generate large, high-dimensional imaging datasets that are challenging to visualize efficiently, particularly at whole-slide and cohort scale. Visualization is an essential step for rapid detection of staining artefacts, such as protein aggregates or non-specific staining. Here, we present Odon, a native Rust desktop viewer designed for rapid, interactive exploration of multiplex imaging data on a standard laptop. Odon is primarily built around the OME-Zarr imaging format, and supports annotations via GeoJSON and GeoParquet, with secondary support for SpatialData, Xenium containers, and TIFF. Data can be stored locally or streamed directly from HTTP or S3-compatible object storage using viewport-driven tile loading. Odon incorporates a highly optimized rendering engine that substantially outperforms existing viewers. In benchmarking, Odon loaded a 32 GB, 36-plex whole-slide OME-Zarr image in under 1 second, compared with 10.14 seconds for QuPath and 35 seconds for Napari. Its GPU-based compositing pipeline also enables smooth rendering and interaction with more than 1,000,000 segmented cells, exceeding the practical limits of many existing tools. Odon further supports integrated visual analytics, including live thresholding and cell selection, and a mosaic mode for simultaneous viewing of hundreds of regions of interest in cohort and tissue microarray studies. Together, these features establish Odon as a high-performance platform for scalable visualization of spatial proteomics data.

## Introduction

Spatial proteomics, a powerful method for the investigation of tissue architecture and tumour microenvironments (Wu et al., 2024; Xu et al., 2024), poses a unique data challenge. Multiplexed platforms such as Akoya PhenoCycler (Goltsev et al., 2018) and the Miltenyi Biotec MACSima (Kinkhabwala et al., 2022) can produce up to 100 high-resolution images of the same tissue section at subcellular resolution, often spanning entire slides, which when scaled across a whole cohort can easily stretch into many terabytes of data. Visual inspection of this data remains a core step for both quality control and analysis. Image stitching errors, antibody aggregates and suboptimal staining are all easily detected by eye. To accommodate this workflow and the sheer size of the data, many labs rely on using a powerful workstation with heavy technical specifications. For example, the technical specifications for MACS iQ View, the Miltenyi Biotec software developed for the MACSima, recommend a 12-core processor (e.g. an Intel Xeon W-3233), RAM of 256 GB or higher, 16 TB of SSD drive storage and an NVIDIA 4060 Ti 8GB variant (“MACS® iQ View - Spatial Biology Software | Miltenyi Biotec | Great Britain,” n.d.). Given that a single-lab or institute will often only have one machine, this leads to a bottleneck in the analysis of the data: only one person can use the machine at a time, whereas a large lab group may have multiple simultaneous projects requiring the machine.

In this study, we leverage two design principles to solve this problem, creating an ultra-fast viewer for spatial proteomics, capable of rendering images of hundreds of samples simultaneously in the same viewer on standard consumer hardware. Firstly, our viewer is constructed using OME-Zarr (Moore et al., 2021) as a core file format, which is a next-generation, cloud-ready file format for biomedical imaging data, and has been demonstrated to outperform TIFF in terms of chunk retrieval time both locally and remotely (either via S3 or HTTP) (Moore et al., 2021), likely due to decreased reliance on a single file handle. Secondly, we employ the state-of-the-art systems programming capabilities provided by the Rust language, which consistently outperforms slower languages like Java and Python (Pereira et al., 2017), which are used for popular viewers QuPath (Bankhead et al., 2017) and Napari (Chiu et al., 2022) respectively. By pairing the lazy-loading capabilities of the OME-Zarr format with Rust’s memory efficiency and robust multi-threading, we have engineered a lightweight solution capable of rapidly rendering high-plex marker panels, whole-slide images and tissue microarrays. This optimized architecture drastically lowers the barrier to entry for spatial data visualization, allowing researchers to perform essential quality control and visual analyses on standard laptops rather than relying on heavily bottlenecked, specialized workstations. Odon derives its name from the dragonfly (*Odonata*), which are reported to exhibit ultra-fast sensitivity to flickering light (Ruck, 1961).

## Results

Odon is a native Rust desktop viewer built to handle the increasing scale of multiplex imaging and spatial omics data. The application supports a wide array of formats, primarily OME-Zarr, GeoJSON, GeoParquet, alongside secondary support for SpatialData, Xenium containers and TIFF (Figure 1a). To bypass local storage bottlenecks, the architecture supports direct remote streaming from HTTP or S3-compatible object storage, utilizing viewport-driven tile loading rather than full dataset downloads.

**Fig. 1.**
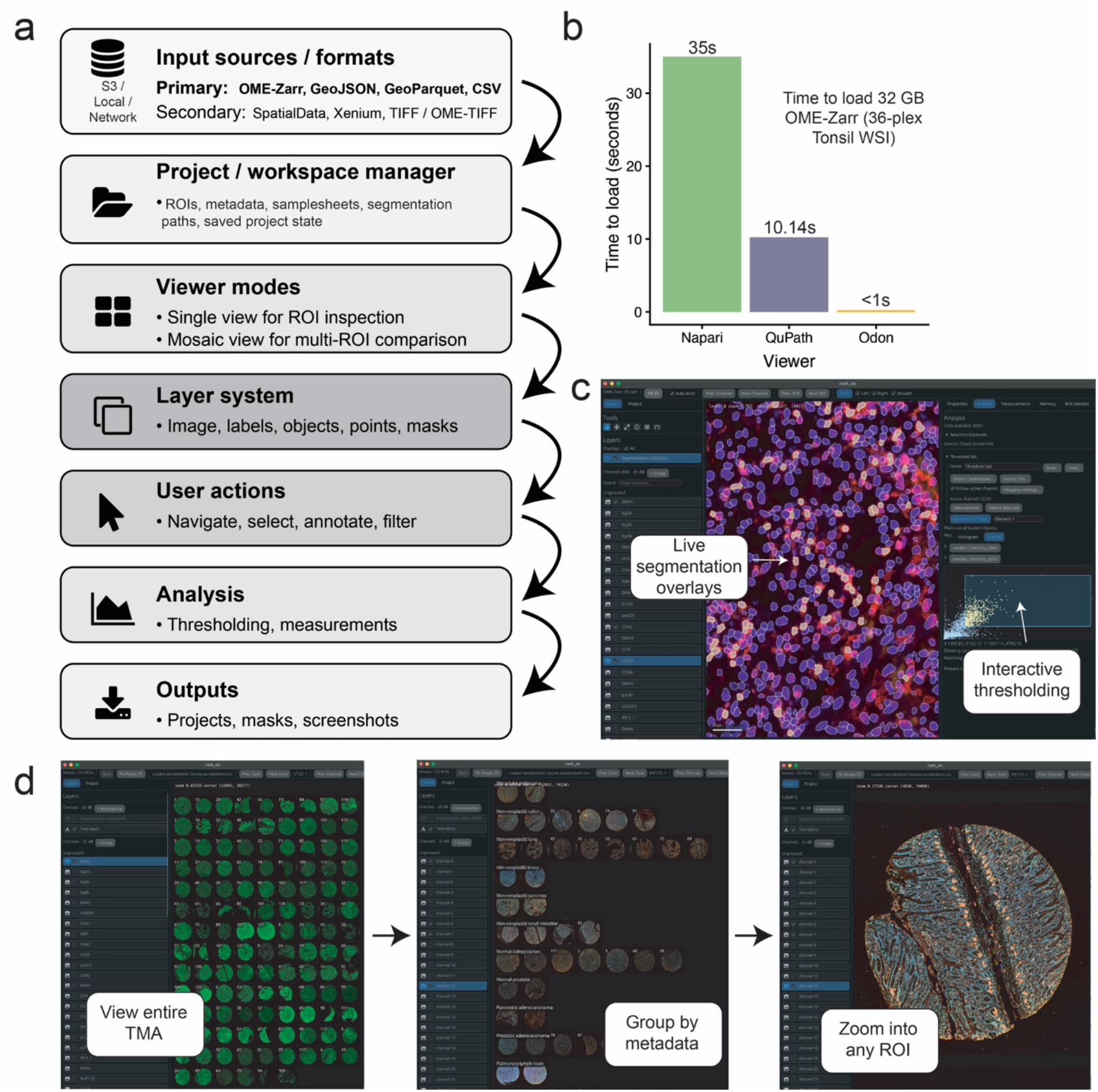
Overview of Odon features and workflow. (a) Schematic of typical Odon workflow. (b) Benchmarking Odon against other popular viewers for spatial proteomics, including Napari and QuPath. The dataset used is the TNP_pilot_cycif dataset from MCMICRO (see data availability). (c) Preview of the live segmentation overlay and interactive thresholding features of Odon. (d) Preview of Odon’s mosaic mode, where hundreds of OME-Zarr datasets, e.g. one from each core of a tissue microarray, can be viewed simultaneously.

A primary advantage of Odon is its highly optimized rendering engine. When benchmarked against widely used platforms, Odon loaded a 32 GB, 36-plex OME-Zarr whole-slide image in under one second, significantly outperforming both QuPath (10.14s) and Napari (35s) (Figure 1b). In addition, when zooming into this image, QuPath caused a system crash. Beyond initial image loading, this performance extends to object rendering via a GPU-based compositing path. While traditional spatial viewers frequently experience severe lag or fail when loading dense segmentation data (often plateauing around 50,000 objects), Odon smoothly renders and interacts with over 1,000,000 segmented cells without performance degradation.

The viewer facilitates granular exploration through a unified layer system supporting images, labels, and masks. Users can seamlessly overlay high-density segmentation maps and interrogate single-cell data in real time (Figure 1c). Odon integrates analysis tools directly into the viewer, including object-property histograms and scatter plots that drive live, on-canvas thresholding and cell selection.

To address the needs of cohort-level studies and tissue microarrays (TMAs), Odon includes a dedicated mosaic mode that displays multiple regions of interest (ROIs) on a single canvas (Figure 1d). This environment is driven by project workspaces or sample sheets, allowing users to automatically group, sort, and label hundreds of tissue cores based on associated metadata. Users can evaluate marker expression across an entire cohort simultaneously, and immediately zoom into any specific ROI for high-resolution inspection.

## Methods

Benchmarking was performed on a 2024 MacBook Pro with 16 GB RAM and a 1 TB SSD hard drive, with the Apple M4 chip, on macOS Sequoia 15.5. To test viewer loading times, the TNP_pilot_cycif dataset from MCMICRO (Schapiro et al., 2021) was used (see data availability), which contains a whole-slide image of a tonsil with 36 channels.

We converted the .tif image to an OME-Zarr v2 multiscale dataset with dimensions 36 x 27,299 x 20,045 (c, y, x) at full resolution. The image pyramid comprised seven levels generated by successive 2-fold downsampling in the spatial dimensions. Image data were stored as 16-bit unsigned integers in 1 x 512 x 512 chunks and compressed using Blosc with the Zstandard codec. The dataset was generated in Python using the zarr and numcodecs packages, with lower-resolution levels derived from the full-resolution image by stride-2 subsampling in X and Y.

For benchmarking against other viewers, we used QuPath version 0.7.0 and napari version 0.6.6 in Python 3.11.13. Load time was measured as a warm start, meaning we did not include the time taken for each respective application to start. We produced screen recordings of each viewer and measured the time until the image completely loaded by examining the length of each video, from image load initiation to the representation of the image on screen, in Adobe Premiere. To test viewing of an entire tissue microarray at once, we used the TMA from the MCMICRO pipeline from the Synapse repository, specifically TMA11, which contains 123 samples. Again, we converted these to OME-Zarr before testing.

## Data availability

Both datasets used for benchmarking are available from the Synapse repository, the TNP_pilot_cycif dataset at https://www.synapse.org/Synapse:syn24849819/wiki/608441 and the TMA data at https://www.synapse.org/Synapse:syn22345748 (TMA11).

## Code availability

Odon is available at https://github.com/alexcoulton/odon

## Acknowledgements

This work was delivered as part of the NexTGen team supported by the Cancer Grand Challenges partnership funded by Cancer Research UK (CGCATF-2021/100014) and the National Cancer Institute (CA278730-01) and The Mark Foundation for Cancer Research. We would also like to thank Dr Karin Straathof, Shreya Yadav, Carmen Rodriguez and Amy Walker for their insightful discussions around spatial proteomics data.

